# Substrate selectivity of the PRDM9 lysine methyltransferase domain

**DOI:** 10.1101/2022.10.12.511945

**Authors:** Jocelyne N. Hanquier, Kenidi Sanders, Christine A. Berryhill, Firoj K. Sahoo, Andy Hudmon, Jonah Z. Vilseck, Evan M. Cornett

## Abstract

Lysine methylation is a dynamic, post-translational mark that regulates the function of histone and non-histone proteins. Many of the enzymes that mediate lysine methylation, known as lysine methyltransferases (KMTs), were originally identified to modify histone proteins but have also been discovered to methylate non-histone proteins. In this work, we investigate the substrate selectivity of the lysine methyltransferase PRDM9 to identify both potential histone and non-histone substrates. Though normally expressed in germ cells, PRDM9 is significantly upregulated across many cancer types. The methyltransferase activity of PRDM9 is essential for double-strand break formation during meiotic recombination. PRDM9 has been reported to methylate histone H3 at lysine residues 4 and 36; however, PRDM9 KMT activity had not previously been evaluated on non-histone proteins. Using lysine-oriented peptide (K-OPL) libraries to screen potential substrates of PRDM9, we determined that PRDM9 preferentially methylates peptide sequences not found in any histone protein. We confirmed PRDM9 selectivity through *in vitro* KMT reactions using peptides with substitutions at critical positions. A multisite λ-dynamics computational analysis provided a structural rationale for the observed PRDM9 selectivity. The substrate selectivity profile was then used to identify putative non-histone substrates, which were tested by peptide spot array. Finally, PRDM9 methylation non-histone substrates were validated at the protein level by *in vitro* KMT assays on recombinant proteins. The selectivity profile of PRDM9 will be useful in identifying putative PRDM9 substrates in different cellular contexts, and future studies are required to determine whether PRDM9 methylates non-histone proteins in the context of meiotic recombination or cancer.

## Introduction

Dynamic lysine methylation, added by lysine methyltransferases (KMTs) and removed by lysine demethylases (KDMs), regulates the function of both histone and non-histone proteins (1, 2). Many KMTs have been studied in the context of histone lysine methylation, which is well understood to regulate chromatin-templated processes. However, many of the KMTs initially identified to modify histone proteins have since been discovered to also methylate non-histone protein substrates (1, 2). Furthermore, several studies have revealed that a large portion of the human proteome is modified with lysine methylation (3–5). To connect lysine methyltransferases with their substrates, we previously developed a functional proteomics approach to map lysine methyltransferase substrate selectivity using lysine-oriented peptide libraries (K-OPL) (6). In this study, we use this approach to characterize the substrate selectivity of the lysine methyltransferase domain of PRDM9.

PRDM9 is a member of the PR-domain containing family of lysine methyltransferases that contain a PR-domain coupled with an array of C2H2 zinc fingers (7–9). The PR-domain is closely related to the SET domain (10), conferring methyltransferase activity, while the zinc fingers allow PRDM proteins to bind to DNA in a sequence-specific manner (11). PRDM9 was first identified as a sterility factor in mice and humans (12, 13). The DNA-binding activity of PRDM9 coincides with hotspots of meiotic recombination (14), and PRDM9 methyltransferase activity is essential for double-strand break formation at PRDM9 designated recombination sites (15). Initial characterization of PRDM9 found that it is typically expressed in female ovaries during development but was only detected in adult testis (13). More recently, an analysis of human patient cancer samples revealed a significant upregulation of PRDM9 across many cancer types (16). However, understanding the role of PRMD9 in these cancer types has been limited by incomplete knowledge of PRDM9 substrates.

The methyltransferase activity of PRDM9 toward histone proteins has been extensively characterized. PRDM9 was first reported to tri-methylate H3K4 (H3K4me3). However, this initial characterization relied upon *in vitro* methyltransferase assays using mouse Prdm9 (herein referred to as mPrdm9; human referred to as PRDM9) and histone substrates isolated from calf thymus and transient overexpression of mPrdm9 in COS-7 cells, with site-specific antibodies used for a limited number of modifications on histone H3 (13). Later, more in-depth studies, including *in vitro* KMT assays followed by mass spectrometry, revealed that mPrdm9 is also capable of H3K4 mono-methylation (H3K4me1) and di-methylation (H3K4me2) (17–19). Experiments using histone peptide arrays also implicated H3K36 as a potential substrate, which was later validated by Eram et al. using *in vitro* reactions on both peptides and nucleosomes (18). Additionally, overexpression of PRDM9 in HEK293 cells increased the global levels of both H3K4me3 and H3K36me3 *in vivo* (18). Another study has suggested that PRDM9 may have substrates on other histone proteins and showed that mPrdm9 mediated the methylation of all four core histone units-H3, H4, H2A, and H2b – alone and within the histone octamer (17). To our knowledge, PRDM9 activity has not been evaluated on non-histone proteins before. In this work, we investigate the substrate selectivity of PRDM9, and one of our key findings is that PRDM9 prefers to methylate peptide sequences not found in any histone protein.

## Results and Discussion

### PRDM9 prefers sequence motifs not found in any histone protein

To determine the preferred sequence determinants of PRDM9, we screened a lysine-oriented peptide library (K-OPL) to query ∼64 million unique peptide sequences subset into 114 sets. Each set contained a fixed central lysine residue (P0) with an additional fixed amino acid within three positions of the central lysine. Transfer of tritium-labeled methyl groups from S-adenosylmethionine to the biotin-labeled K-OPL peptides was detected using a sensitive surface proximity assay (6). Three independent reactions for each of the 114 separate sets were averaged and normalized to the highest signal across all sets for each amino acid fixed at a particular position relative to the central lysine residue (Fig. 1A). The selectivity analysis reveals that PRDM9 prefers isoleucine (Ile; I) in the P-1 position. To confirm the signal of PRDM9 on the K-OPL peptides, we selected several K-OPL sets with high, medium, or low signal (see methods) at three different positions and performed additional methyltransferase assays confirming the preference for each amino acid at these positions (Fig. S1C). To check the potential impact of neighboring post-translational modifications on PRDM9 activity, we also performed analysis of additional K-OPL sets that contained phosphorylated serine (Ser; S), tyrosine (Tyr; Y), or threonine (Thr; T). PRDM9 showed a preference for Ser in the P-3 position, and Thr in the P+2 position. Phosphorylation of these residues reduced activity to near background levels (Fig. S1D), highlighting the potential for post-translational modification crosstalk (1).

**Figure 1.**
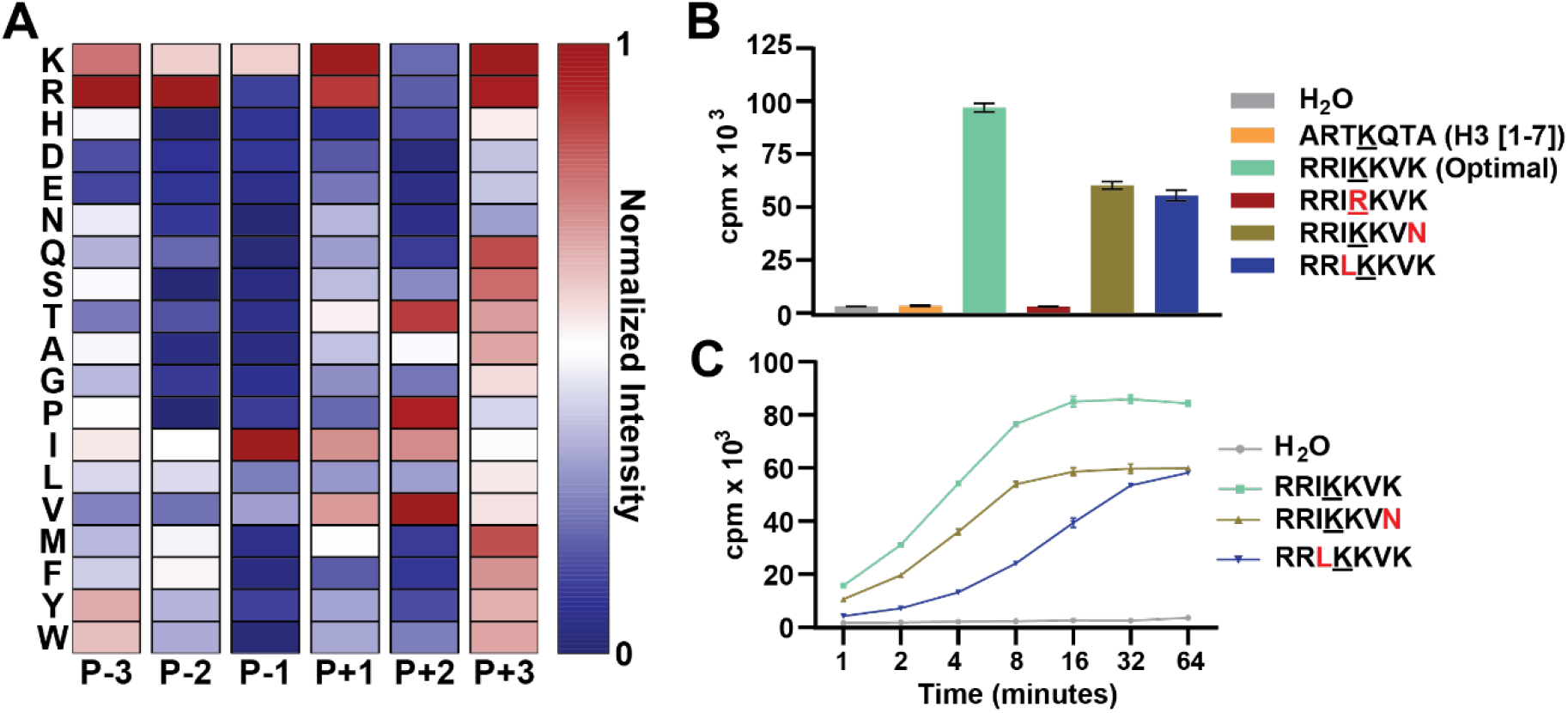
Substrate selectivity of PRDM9. *A*, K-OPL substrate selectivity profile for PRDM9. Averaged results from three independent K-OPL surface proximity assay screens for PRDM9 depicted as a position-normalized heat map (see Fig. S1 for globally normalized heat map and raw K-OPL data); the exception is duplicate measurements for P+2 M. The color code is proportional to the creation of enzyme product, where red (1) is most active and blue (0) is least active. Rows show the identity of each fixed residue, and columns show the position within the sequence. *B*, Validation of PRDM9 substrate selectivity using peptides with substitutions to critical residues. KMT reactions consisted of 0.8 µg PRDM9, 0.5 µg substrate, and 2 µCi SAM, and were incubated for 1 hr at room temperature. Graph displays mean (n=3) ±SD. Cpm = counts per minute. *C*, Initial rate measurements with the optimal PRDM9 substrate (RRI<u>K</u>KVK, bright green) and peptides containing substitutions at P-1 (I to L, blue) and P+3 (K to N, moss green) positions. KMT reactions consisted of 0.8 µg PRDM9, 0.5 µg substrate, and 2 µCi SAM. Graph displays mean (n=3) ±SD; error bars are masked by the symbol weight for some data points.

To validate the PRDM9 substrate selectivity profile, a series of peptides were synthesized based on the optimal substrate (RRI<u>K</u>KVK). PRDM9 methylated the optimal substrate nearly 100-fold more efficiently than a peptide containing H3K4 (Fig. 1B). As predicted, the substitution of the P-1 Ile with leucine (Leu; L) resulted in a significant reduction in PRDM9 methyltransferase activity (Fig. 1, B and C). Importantly, there was no signal when using a peptide with the central lysine (Lys; K) substituted with arginine (Arg; R) (Fig. 1B). The position normalized and globally normalized heatmaps of PRDM9 substrate selectivity (Fig. 1A; S1B) suggest PRDM9 has a stronger preference for the P-1 position than P+3. To evaluate this, we also tested a peptide with lysine substituted with asparagine (Asn; N) in the P+3 position, which resulted in reduced activity but had less impact than the substitution made in the P-1 position (Fig. 1, B and C).

Next, we evaluated how substitutions surrounding the previously reported PRDM9 substrate H3K4 impacted PRDM9 activity. A series of H3 peptides were synthesized with the native P-1 Thr substituted with Ile, Leu, or valine (Val;V). *In vitro* methyltransferase assays using these peptides as substrates recapitulated the rank order observed in the K-OPL screen for these residues in the P-1 position (Fig. 2A). The Ile peptide was methylated most efficiently, followed by Val, Leu, and then Thr. Similar analysis on a peptide with the natural P+1 Glu substituted with a Lys further confirmed the preferences identified by the K-OPL substrate screen. A peptide containing Lys at the P+1 position in the context of an H3K4 peptide resulted in a significant increase in PRDM9 activity. Overall, these results further confirm the accuracy and predictive power of the K-OPL selectivity profile for PRDM9 substrates.

**Figure 2.**
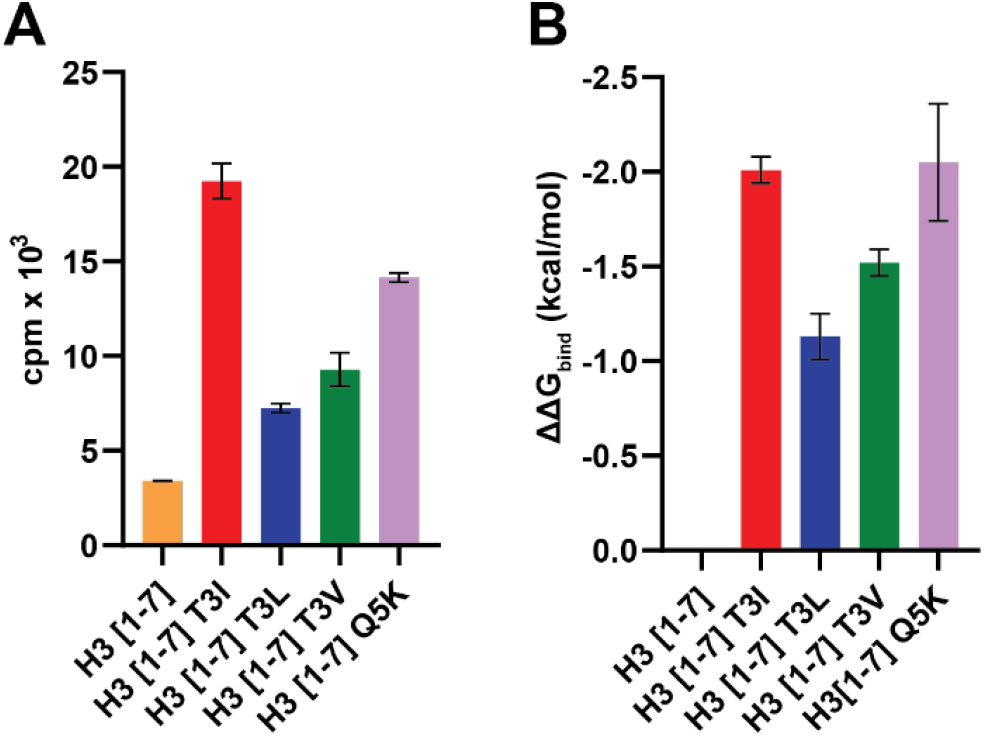
MSλD analysis confirms PRDM9 substrate selectivity. Comparison of *in vitro* lysine methyltransferase assays on histone H3 peptides (A) and relative changes in binding free energies (ΔΔ*G*bind) calculated from MSλD simulations (B). Bar graphs display mean ±SD (n=3 for A and n=5 for B).

### Multisite λ-dynamics (MSλD) analysis identifies a structural rationale for PRDM9 substrate selectivity

To determine the molecular basis for PRDM9 substrate selectivity, a computational and structural analysis of PRDM9-peptide binding was performed with multisite λ-dynamics (MSλD). MSλD is a physics-based alchemical free energy method that can provide structural insights into molecular binding via molecular dynamics sampling and can quantify the thermodynamic effects of a protein side chain mutation on peptide-protein binding (20–22). Using a previously reported mPrdm9 SET-domain structure solved in complex with an H3K4me2 peptide and S-adenosylhomocysteine (19), four peptide-substrate substitutions were investigated with MSλD. It is important to note that the mPrdm9 has a high sequence identity to human PRDM9 (72.8%). Furthermore, we compared the substrate selectivity of hPRDM9 and mPrdm9 on the same H3K4 peptides used in MSλD analysis and found no differences (Fig. S2A).

At the P-1 position, the native Thr was mutated to Ile, Leu, and Val; then, at the P+1 position, the native glutamine (Gln;Q) was mutated to Lys. Computationally, the experimental K-OPL activity trends (Ile > Val > Leu > Thr at P-1; Lys > Gln at P+1) were reproduced (Fig. 2A). Additional targeted experiments confirmed these predictions and demonstrated a strong positive correlation between predicted binding affinities of different peptide mutants and their observed methylation activities by PRDM9 (Pearson’s r = 0.887; Fig. S2B). These results suggest a strong correlation between binding affinity for PRDM9 peptide substrates and enzymatic activity. This work also demonstrates that the MSλD analysis is effective for identifying and screening preferred peptide substrates for PRDM9 and may be beneficial for screening other KMTs.

MSλD trajectories were analyzed to determine the molecular features that drive PRDM9 preference for Ile in the P-1 position and Lys in the P+1 position. A comparison of the distance between the ε-amine of the substrate lysine and the S-adenosylmethionine showed no significant changes (Fig. S3A), suggesting no direct influence on the catalytic proficiency of the enzyme by the different substitutions. Rather, an induced fit model of complementary size, shape, and nonbonded interactions was observed. Analysis of the Cα-Cβ dihedral angles at the P-1 position indicated that each residue had slightly different rotational preferences when bound to PRDM9. Thr adopts two primary conformations to form a hydrogen bond with the backbone carbonyl group of Prdm9 Ala287 or, when rotated ∼85°, to expose the hydroxyl group to solvent (Fig. 3, A, B, and C). In either conformation, Thr suffers a slight desolvation penalty for binding PRDM9, since it is partially shielded by PRDM9 and can form only 1-2 hydrogens bonds at a time (approximately 1 fewer hydrogen bond than Thr would form in bulk solvent). In contrast, Val, Ile, and Leu show favorable hydrophobic packing against PRDM9 residues Ala287, Tyr304, and Leu294 at the P-1 position (Fig 3, A, D, E, and F). Ile is slightly favored over Val due to its increased size and interactions with these hydrophobic residues, although both Ile and Val branch at Cβ to bind similarly as Thr, and both share similar Cα-Cβ dihedral angle preferences (Fig. 3, D and E). Leu is the least preferred of the hydrophobic residues tested due to minor steric clashes made with Tyr304. This is evident in the longer average distances for Leu between its Cα atom and the Y304 aromatic ring center of mass compared to Thr, Ile, or Val (Fig. S3C). This is largely due to leucine’s branching at the Cγ atom, which alters its conformational preferences in both Cα-Cβ (Fig. 3F) and Cβ-Cγ (Fig. S3B) dihedral angle distributions. Finally, the preference for Lys in the P+1 position is readily explained by the introduction of favorable ionic interactions between the P+1 Lys with residues Glu360 and Asp359 (Fig. 3G). This also explains why Arg would be favorable at the P+1 position, as it will likely form similar favorable ion-ion interactions. Overall, this MSλD analysis provides a structural rationale for PRDM9 substrate selectivity that supports the K-OPL derived substrate selectivity profile and provides mechanistic insight into the striking preference for Ile in the P-1 position and Lys at the P+1 position.

**Figure 3.**
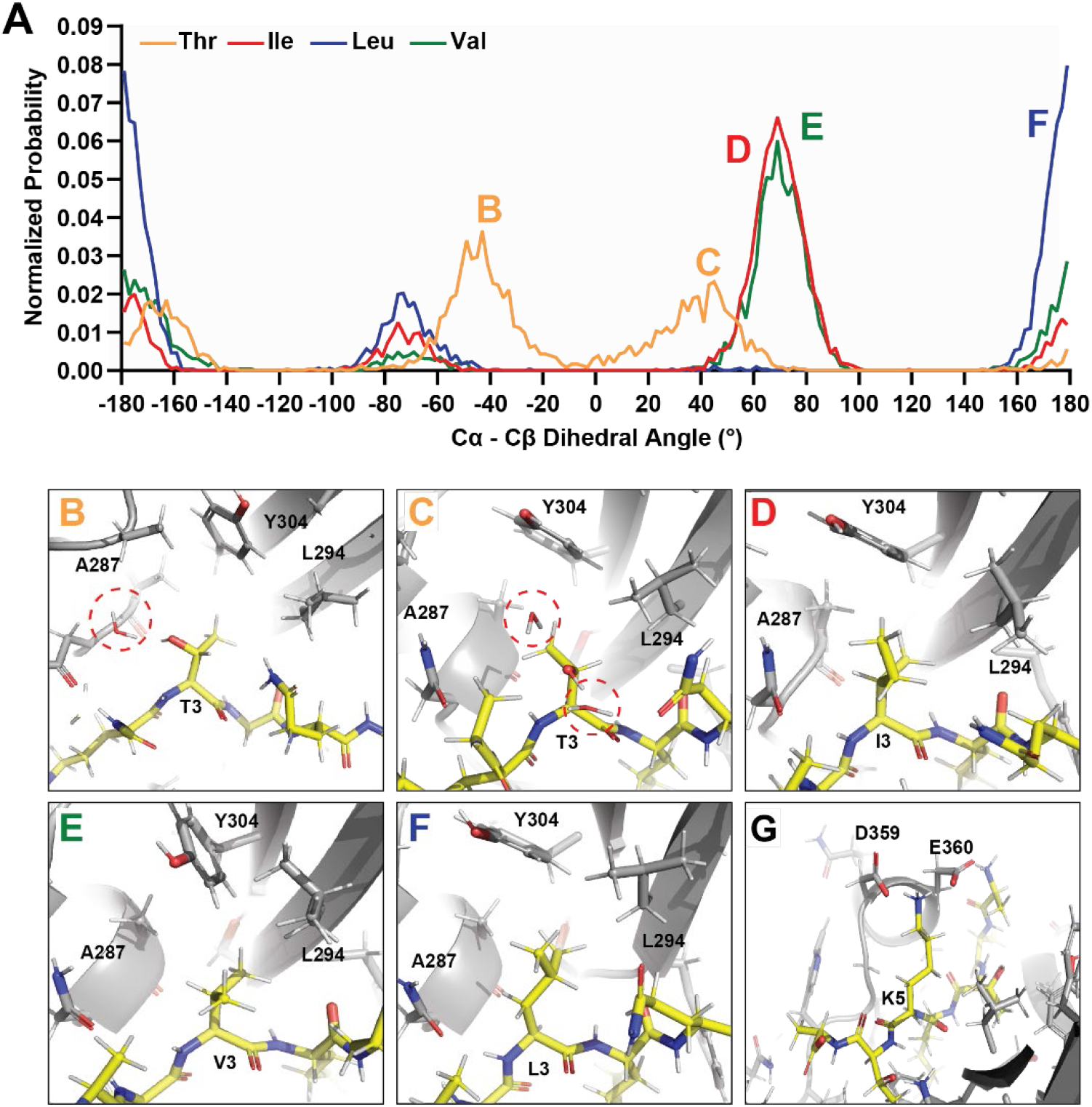
Analysis of MSλD trajectories reveal structural rationale for PRDM9 substrate selectivity. *A*, Distribution of Cα-Cβ dihedrals from MSλD. Representative images from MSλD trajectories for native Thr3 (*B and C*) or substitutions Ile3 (*D*), Val3 (*E*), Leu3 (*F*) or Lys5 (*G*).

### Identification of non-histone peptide substrates of PRDM9

We next identified and evaluated putative PRDM9 substrates based on the selectivity profile: we used the PRDM9 K-OPL selectivity data to score each lysine-centered 7-mer motif found in the human proteome (Fig. 4A and Table S1) and synthesized a peptide spot array to test twenty-five putative PRDM9 substrates. The putative substrates were chosen from the top 15% of all lysine-centered 7-mer motifs found in human proteins. The top 15% represent 137 sequence motifs, which were narrowed down to twenty-five based on manual curation (Table S2). Factors for inclusion included expression in spermatocytes, one of the few tissues where PRDM9 is expressed, connection to double-strand break formation or DNA repair, and/or the predicted methylated lysine residing within an annotated protein domain/region (Table S3). In addition to putative substrates, the peptide array also contained two negative control peptides, peptides containing the known histone substrates of PRDM9 (H3K4 and H3K36), and the optimal PRDM9 peptide. Each peptide was spotted in triplicate, and to determine if other lysine residues in the peptides were being methylated, a corresponding peptide with the predicted target lysine substituted with an arginine was also included. PRDM9 methylated eighteen of the twenty-five putative substrates, and twelve substrates showed more signal than either histone peptide control (Fig. 3B). All seven putative substrates that were not methylated did not contain an Ile in the P-1 position but had nearly optimal sequences in all other positions, further highlighting the importance of the P-1 Ile for PRDM9 substrate selectivity. PRDM9 methylated the H3K36 peptide more efficiently than H3K4, likely due to the P-1 Val in H3K36, which the K-OPL selectivity profile shows is preferred over the Thr found at this position in the context of H3K4. Substitution of the Lys predicted to be methylated with Arg resulted in a complete loss of any detectable methylation signal for nearly all peptides. A notable exception is ACE K722, which showed a significant signal even in the Lys-to-Arg control but also contained an additional Ile-Lys motif.

**Figure 4.**
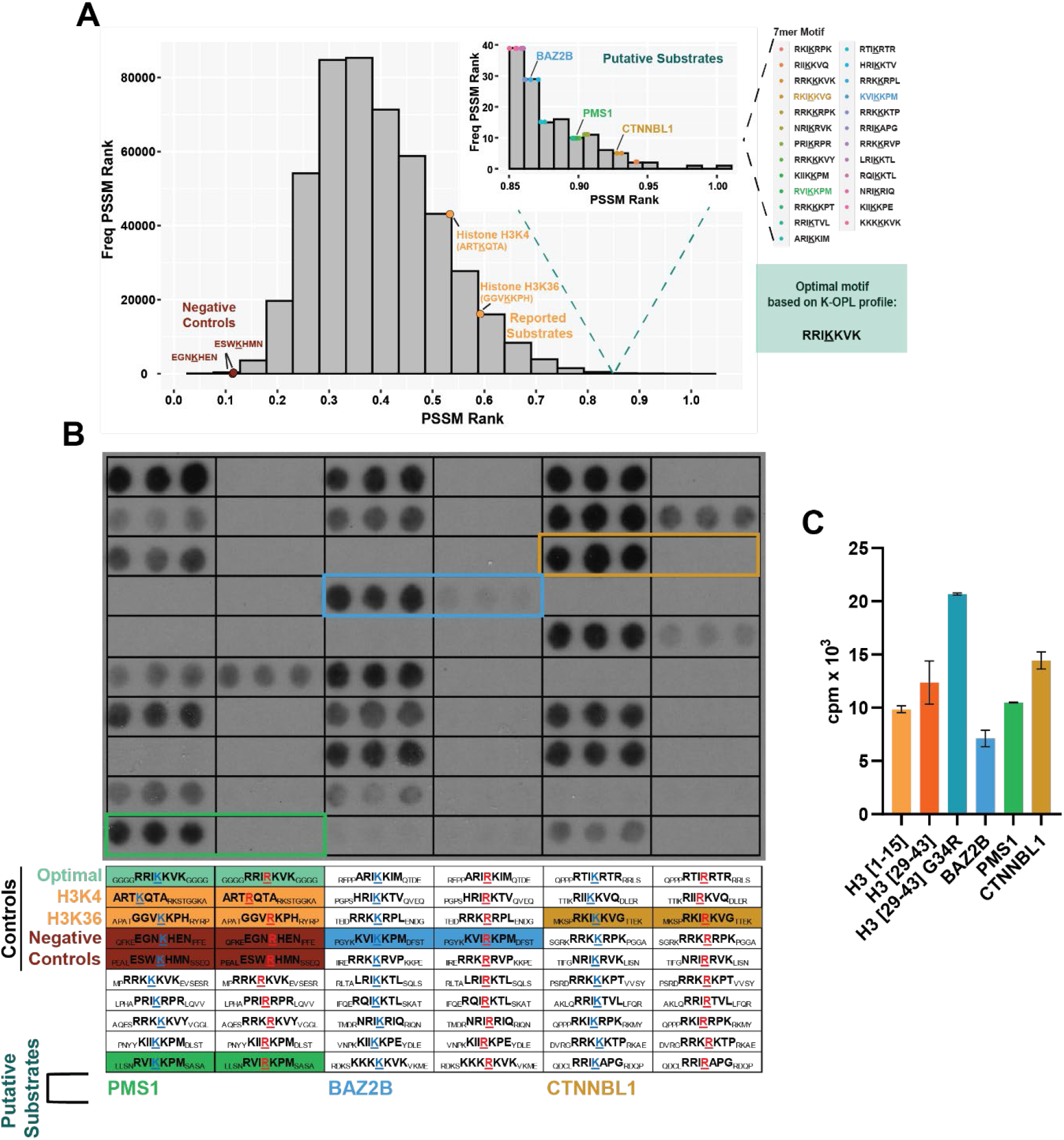
Methylation of non-histone protein derived peptide substrates by PRDM9. *A*, Distribution of position-specific scoring matrix (PSSM) scores for all 7-mer motifs surrounding a central lysine in the human proteome based on PRDM9 signal on K-OPL sets. Substrates used for spot array are highlighted, including: negative controls with low PSSM scores (maroon), reported histone substrates (orange), and putative substrates with PSSM scores ranked in the top 15% of all scores (inset). *B*, Autoradiography signal from PRDM9 lysine methyltransferase activity on peptide spot array (15-mer peptides spotted in triplicate). One hour film exposure shown. The location and sequence of each feature are displayed below. *C*, Comparison of newly identified non-histone substrates with histone peptides. Graph displays mean (n=2) ±SD.

We selected three substrates for further study – BAZ2B, PMS1, and CTNNBL1 – that PRDM9 robustly methylated and whose predicted methylation site is located within a functional protein domain/region. The lysine residues predicted to be methylated within BAZ2B, PMS1, and CTNNBL1 reside within the bromodomain (BRD), high mobility group (HMG) box, and armadillo (ARM) repeat, respectively (Fig. S4). Lysine methylation in these domains on other proteins has previously been shown to alter protein function, localization, or stability (23–25). To further test whether PRDM9 methylated these proteins, we synthesized peptides corresponding to BAZ2B, PMS1, and CTTNBL1 sequences. PRDM9 methylated all three peptides to a similar degree as histone peptides, with CTNNBL1 showing the highest signal (Fig. 3C); CTNNBL1 was also ranked the highest of these three putative substrates (Fig. 3A, inset) according to the K-OPL selectivity profile. All three substrates contained the P-1 Ile and P+1 Lys validated as critical residues for PRDM9 substrate activity.

In addition to testing these putative substrates, we also synthesized a peptide centered around H3K36 with Gly 34 substituted with Arg, a variant found in glioblastoma and osteosarcoma (26–29). Structural studies of SETD2, another KMT that mediates H3K36 trimethylation (H3K36me3), showed that the bulky Arg substitution prevents substrate engagement (30). As a result, H3G34R was shown to decrease global trimethylated H3K36 on molecules with the substitution (27). However, the PRDM9 K-OPL selectivity profile predicts Arg would be preferred over Gly at this position. Indeed, PRDM9 methylated the H3G34R peptide more efficiently than wild-type H3K36. A recent study demonstrated that PRDM9 is expressed in some glioblastomas, and studies of H3G34R have identified a redistribution of H3K36 methylation with some loci modified with a higher frequency (31). Our data underscore the possibility that other KMTs may have increased activity toward some variant histone proteins, perhaps explaining why some of these histone mutations result in a redistribution of the modification rather than decreases or increases alone (31).

### PRDM9 methylates non-histone proteins

To determine whether PRDM9 methylates BAZB2, PMS1, and CTTNBL1 at the protein level, we cloned and purified each protein. *In vitro* KMT assays were performed using tritiated S-adenosylmethioine with recombinant BAZ2B, PMS1, and CTNNBL1 as putative substrates and histone H3 as a positive control. The reactions were separated on SDS-PAGE gels, treated with an autoradiography enhancer, dried, and exposed to film to detect methylated proteins by autoradiography. As expected, PRDM9 methylated histone H3; PRDM9 also methylated PMS1 and CTNNBL1 (Fig. 5A). Only a faint band was detected for BAZ2B. Analysis of available structural data for BAZ2B shows the predicted methylation motif is in an alpha-helix (PDB: 3G0L), which may explain why BAZB2 is not methylated more efficiently by PRDM9. Because PRDM9 displayed marked lysine methyltransferase activity on CTNNBL1, we sought to validate that the autoradiography signal on CTNNBL1 was due to methylation of the lysine predicted to be methylated, K394. We performed *in vitro* KMT assays using recombinant CTNNBL1 with and without the target lysine substituted with arginine (K394R) (Fig. 5B). PRDM9 methyltransferase activity on wild-type CTNNBL1 is evident; in contrast, there was no detectable activity on CTNNBL1 K394R, supporting the conclusion that PRDM9 methylates CTNNBL1 at K394. When comparing the activity of PRDM9 on CTNNBL1 to that on histone H3, it is important to note that PRDM9 methylates CTNNBL1 at one residue and methylates histone H3 at multiple residues, which could contribute to the higher autoradiography signal on histone H3.

**Figure 5.**
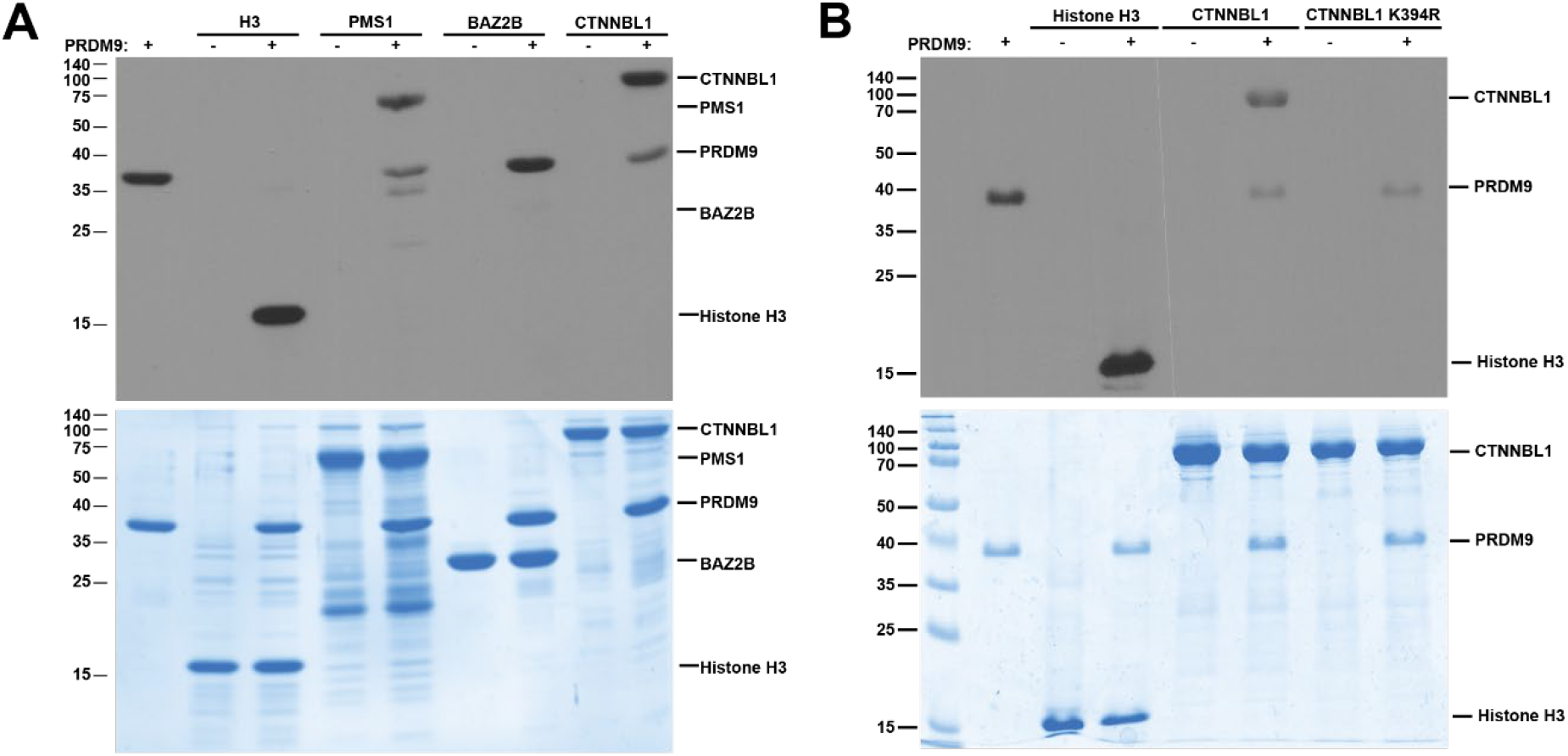
PRDM9 methylates non-histone proteins *in vitro*. *A*, PRDM9 activity on recombinant non-histone substrates detected by fluorography. Representative of two independent experiments. *B*, Comparison of PRDM9 methylation of CTNNBL1 when the target Lys is substituted with and Arg (K394R). Representative of three independent experiments. Top panels depict fluorography signal, and the bottom panel shows total protein stained with Coomassie.

## Conclusions

The data presented in this study strongly suggest that PRDM9 has physiologically relevant non-histone substrates. *In vitro* K-OPL screening revealed that PRDM9 prefers to methylate a sequence motif not found in any histone protein, and structural rationale for PRDM9 selectivity was provided by multisite λ-dynamics computational analysis. Furthermore, we have demonstrated that PRDM9 methylates recombinant CTNNBL1 protein at K394. Lysine 394 resides within an ARM repeat region of CTNNBL1; ARM repeats form an α-superhelix, providing a platform for protein interactions, including activation or degradation of the target protein (32). In the case of the related protein β-catenin, regulation of lysine methylation within an ARM repeat region was shown to regulate protein stability (24). Future studies are necessary to determine if PRDM9 methylates CTNNBL1 in cells and whether there are functional impacts. Overall, the PRDM9 K-OPL selectivity profile will be helpful in identifying whether non-histone substrates contribute to the regulation of recombination or play a role in the cancers in which PRDM9 is overexpressed, and future studies are necessary to determine whether PRDM9 methylates non-histone proteins in these contexts.

### Experimental Procedures

#### Protein expression and purification

Recombinant human PRDM9 (amino acids 191-415) with an N-terminal GST tag (Active Motif) was used for experiments shown in Figures 1 and S1; all other experiments were performed using recombinant PRDM9 purified as follows. Human PRDM9 (amino acids 195 to 385), mouse Prdm9 (amino acids 198-368), BAZ2B (amino acids 2062-2166), PMS1 (amino acids 333-705), and CTNNBL1 (full length, amino acids 1-563) were cloned in a pET28b expression vector as 6XHis-SUMO N-terminal fusions. Point mutations were generated by QuikChange Site-Directed Mutagenesis (Stratagene). Constructs were transformed into BL21(DE3) cells and plated on LB agar plates containing kanamycin. A single colony was selected to grow a starter culture in LB media with kanamycin overnight at 37°C. The starter culture was diluted 100-fold into 1L of LB media in a 2L baffled shaker flask and grown at 37°C shaking at 160rpm until the OD600 (optical density at 600 nm) reached 0.6 to 0.8, at which point the temperature was lowered to 16°C, isopropyl-β-D-thiogalactopyranoside was added (0.5 mM), and incubation was continued overnight with shaking at 160rpm. Bacteria were harvested by centrifugation and either frozen at −80°C or used immediately. Cells were resuspended in lysis buffer (20 mM Tris pH 8.0, 500 mM NaCl, 5 mM Imidazole, 1mM DTT, 1mM PMSF) and lysed by passing through a microfluidizer. Cell lysates were cleared by centrifugation at 14,000 x g for 30 minutes. After incubating cleared lysate with Pierce Ni-IMAC resin for 1 hour at 4°C, the resin was washed with lysis buffer and bound protein was eluted using elution buffer (20 mM Tris pH 8.0, 500 mM NaCl, 500 mM Imidazole, 1 mM DTT, 1mM PMSF). Eluted protein was concentrated and buffer exchanged using a 10K MWCO Amicon Ultra Centrifugation device, followed by size exclusion chromatography (SEC) on a Cytiva Superdex 200 Increase column using an AKTA Pure FPLC and SEC buffer (50 mM Tris pH 8.0, 150 mM NaCl). Fractions were separated by SDS-PAGE and stained with Coomassie. Fractions containing the predicted proteins were concentrated using an Amicon Ultra Centrifugation device, aliquoted, and stored at −80°C in SEC buffer.

#### Lysine methyltransferase surface proximity assay (SPA)

Except when otherwise stated, for reactions with K-OPL sets or biotin labeled peptides, reactions (10 μl) containing 0.8 μg of PRDM9, 0.5 μg of a substrate, and 2.2 μCi of ^3^H-SAM (PerkinElmer) in KMT reaction buffer [50 mM tris (pH 8.8), 5 mM MgCl2, and 4 mM dithiothreitol] were incubated for 6 hours at room temperature. Reactions were stopped by adding trifluoracetic acid to a final concentration of 0.5%, neutralized by diluting with 135 μl of 50 mM NaHCO3, and transferred to white 96-well microplates (PerkinElmer). 8 μl of streptavidin-coated SPA bead slurry (0.1 mg/μl, PerkinElmer) was added to each well, the plate was sealed using PerkinElmer TopSeal-A, and centrifuged for 3 minutes at 1000 rpm. The bead slurry was incubated with the reaction mixture for 30 minutes before liquid scintillation counting for 1 minute using a Hidex Sense Beta microplate reader. K-OPL peptides with high, medium, and low signal were selected for positions P-3, P-1, and P+1. In each case, the fixed amino acid K-OPL set with the highest signal was selected as the high signal group. The fixed amino acid K-OPL set with approximately half the signal was selected as the medium example and the low signal was a K-OPL set with signal near the background.

#### Peptide spot assays

Putative PRDM9 substrates were synthesized as 15-mer peptides on a β-alanine derivatized membrane (Amino-PEG500-UC540 membrane, Intavis) using Fmoc (N-(9-fluorenyl) methoxycarbonyl) protected activated amino acids (CEM) and a solid phase synthesis protocol described previously (33, 34). The membranes were deprotected with a deprotection cocktail that includes 88% Trfluroacetic acid, 5% water, 5% Phenol, and 2% Triisopropyl Silane, followed by three washes with dichloromethane, three washes with dimethyl formamide and three washes with ethanol as described previously (33, 34). The deprotection protocol was repeated twice, and the membrane was allowed to dry and stored at −20°C. Putative PRDM9 substrates (15-mer peptides) were synthesized in triplicate, and peptides with the lysine residue predicted to be methylated substituted with arginine were included as a control. To test PRDM9 methylation activity on putative substrates, the peptide array was first pre-incubated in KMT reaction buffer [50 mM tris (pH 8.8), 5 mM MgCl2, and 4 mM dithiothreitol, 0.01% Triton X-100] for 20 minutes; following which, the membrane was incubated in KMT reaction buffer supplemented with ^3^H-SAM (PerkinElmer) and 6xHis-SUMO-tagged human PRDM9 (0.5 µCi and 0.4 µg, respectively, per 10 µL reaction) for 1 hour at room temperature. The membrane was then washed 5 times for 5 minutes in washing buffer (1% SDS in PBS) and incubated in Amplify fluorographic reagent (Cytiva) for 5 minutes. All incubation and washing steps were carried out using a shaker. The membrane was exposed to film at −80 C° in the dark and film was exposed following a range of exposure times as indicated.

#### In vitro lysine methyltransferase reactions

For reactions with protein substrates, 0.125 μg of PRDM9, 1 μg of the indicated substrates, and

1 μCi of ^3^H-SAM (PerkinElmer) in KMT reaction buffer were incubated for 1 hour at room temperature. Purified human histone H3 protein was purchased from Active Motif. Reactions were quenched by the addition of SDS loading buffer and resolved by SDS–polyacrylamide gel electrophoresis. Following the detection of total protein by Coomassie staining, gels were treated with EN3HANCE (PerkinElmer), dried, and methylated proteins were detected by autoradiography.

#### Multisite λ-Dynamics Analysis

A structural rationale for PRDM9’s observed K-OPL binding preferences was investigated computationally with MSλD (20–22). A previously reported *mouse* Prdm9 SET-domain structure (PDBID: 4C1Q) with an H3K4me2 peptide and S-adenosylhomocysteine (AdoHcy) was used to provide a structural model of *holo* human PRDM9 (19) To reverse the catalytic reaction, AdoHcy was converted to S-adenosyl methionine (SAM) and the H3K4me2 peptide was reverted to its mono-methylated form H3K4me1. Potential residue flips and protonation state assignments for titratable groups were determined with the assistance of Molprobity and ProPKa (35–37). The system was then solvated with the CHARMM-GUI webserver (38); all crystallographic solvent molecules were retained and a cubic box of TIP3P water molecules was generated to solvate the peptide-PRDM9 complex (39). A neutralizing buffer of 0.1M of Na^+^Cl^−^ was also added to the simulation cell. A similar setup was followed for preparing a solution of the isolated peptide. All CHARMM-based force fields were used to represent the components of the chemical system, including CHARMM36 for protein and peptide molecules and CGenFF for the SAM cofactor (40–44). MSλD free energy calculations were run in the CHARMM molecular simulation package on graphical processor units (GPUs) with BLaDE (44–46). All simulations were run in the isothermal-isobaric ensemble at 25° C and 1 atm. SHAKE was used to restrain all heavy-atom-Hydrogen bond lengths (45). Long-range interactions were gradually smoothed to zero with force switching from 9 – 10 Å, and particle mesh Ewald was used to correct for long-range electrostatic interactions (47–50). Soft-core non-bonded potentials were used to avoid endpoint singularities for alchemical sampling (51). Prior to MSλD production sampling, each system was subject to 250 steps of steepest descent minimization to remove potential clashes. Biasing potentials for MSλD were then determined with the Adaptive Landscape Flattening algorithm over a combined 173 ns of sampling (51). To determine the final relative free energy differences, five replicate 25 ns MSλD production simulations were performed. In accordance with a standard alchemical thermodynamic cycle for calculating binding affinities, residue mutations were investigated in both isolated peptide and peptide-PRDM9 bound states (19). Additionally, for the Gln to Lys mutation at the P+1 position of the peptide substrate, the alchemical ion approach was used to maintain charge neutrality throughout the duration of a charge-changing MSλD perturbation. The alchemical ion-water pair was restrained with a harmonic force constant of 59.2 kcal/mol·Å^2^ (52, 53). All dihedral and distance analyses were performed with CHARMM, and trajectory analyses and figures were made with PyMOL (54).

#### Bioinformatics analysis

A position specific scoring matrix (PSSM) score for PRDM9 selectivity was calculated for every lysine-centered 7-mer in the human proteome based on PRDM9 activity on K-OPL sets. The input for PSSM score calculation was a normalized and transformed matrix of PRDM9 signal on K-OPL positions P-3 to P+3. The average cpm of three independent measurements was globally normalized to the highest cpm value, a pseudocount (+1) was added to each normalized count, and then the natural log was taken. PSSM scores were calculated on the normalized and transformed matrix by summing the score for each amino acid at each position +/-3 residues from a central lysine.

## Data Availability

All data are included in the manuscript and supporting information.

## Supporting information

This article contains supporting information.

## Author contributions

J.N.H. and E.M.C. designed all experiments and discussed results. J.N.H., K.S., and C.A.B. performed methyltransferase assays under E.M.C.’s supervision. A.H. and F.K.S. designed and fabricated peptide SPOT arrays. J.Z.V. performed the MSλD simulations and computational analysis. J.N.H. and E.M.C wrote the manuscript with input from all authors.

## Funding and additional information

This work was supported by start-up funds from the Indiana University School of Medicine Department of Biochemistry and Molecular Biology, Precision Health Initiative, and the Melvin and Bren Simon Comprehensive Cancer Center to E.M.C and J.Z.V. This work was also supported by National Institutes of Health grant R35GM147023 to E.M.C. The content is solely the responsibility of the authors and does not necessarily represent the official views of the National Institutes of Health or Indiana University. J.Z.V. gratefully acknowledges the Indiana University Pervasive Technology Institute for providing supercomputing and storage resources that have contributed to the research results reported within this paper.

## Abbreviations

The abbreviations used are:

KMT: lysine methyl transferases
K-OPL: lysine-oriented peptide libraries
MSλD: multi-site λ-dynamics
PSSM: position-specific scoring matrix
PTM: post-translational modification.

---

**Figure S1.**
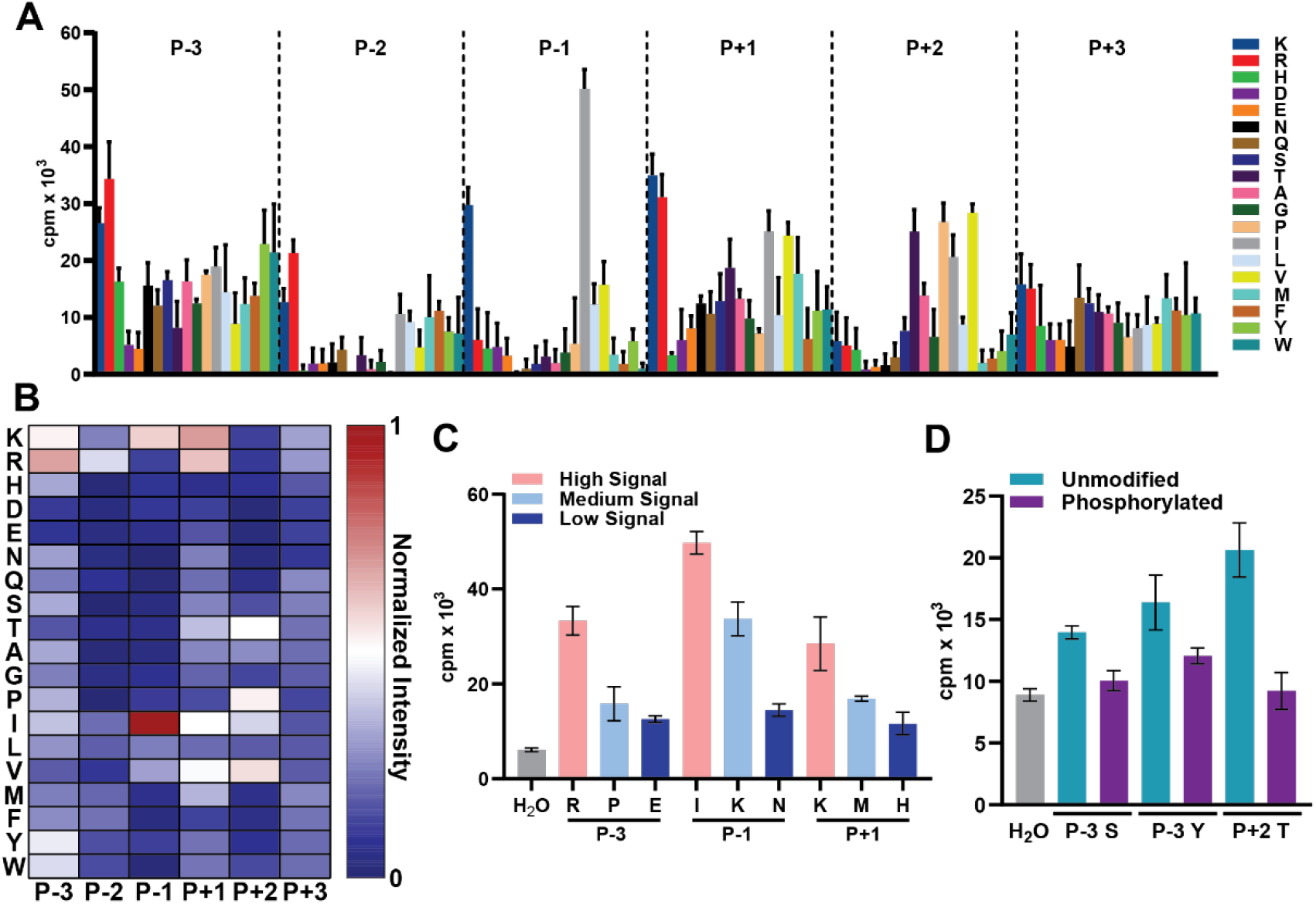
Analysis of PRDM9 substrate selectivity using a K-OPL. *A*, Data for PRDM9 K-OPL selectivity profile showing average signal for reactions with each K-OPL set (n=3). *B*, Heatmap normalized globally to the set with the highest signal. *C*, Validation of PRDM9 K-OPL selectivity profile on select K-OPL sets (n=2). *D*, PRDM9 reactions on phosphorylated K-OPL sets (n=3). Bar graphs display mean ±SD.

**Figure S2.**
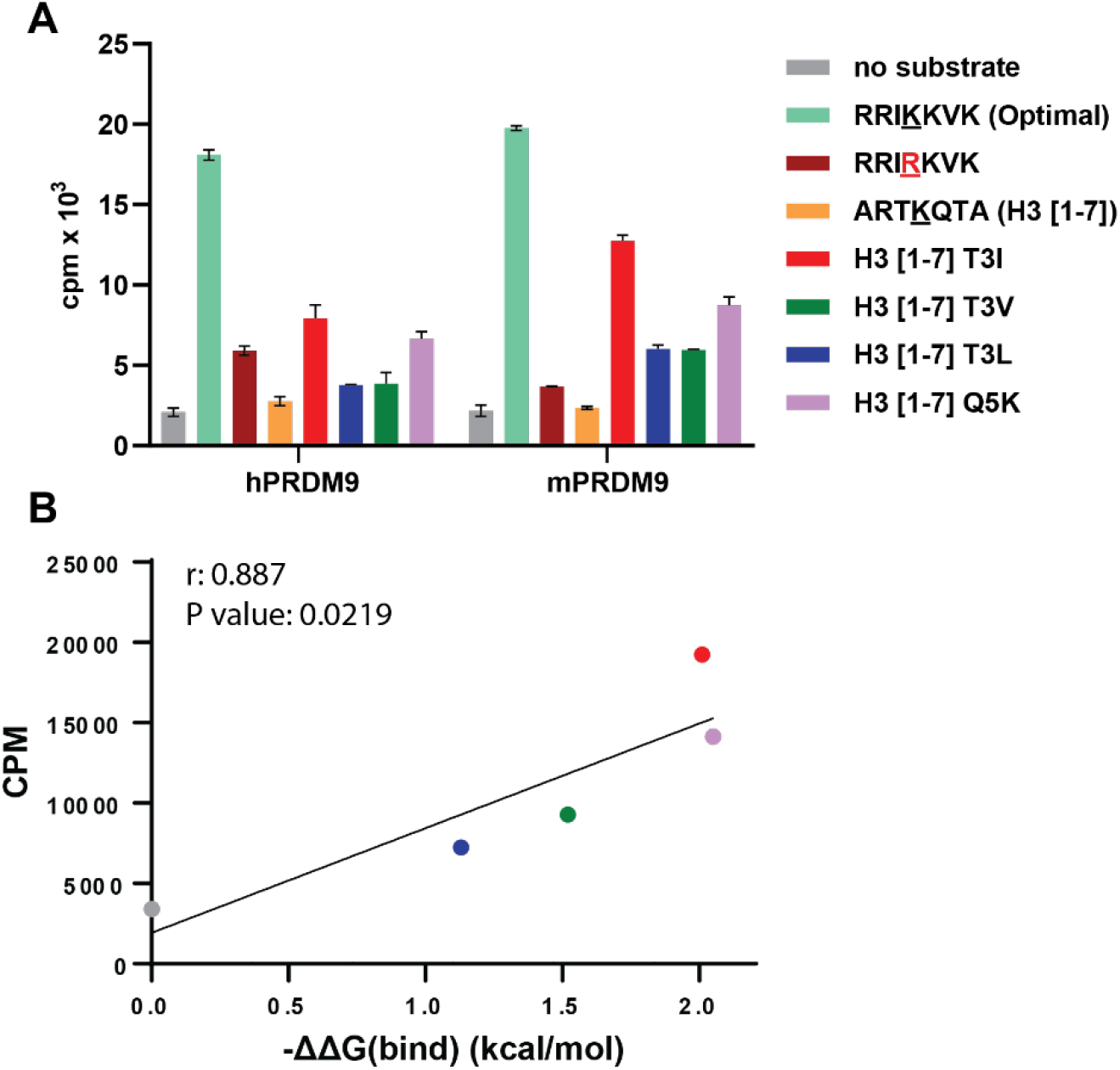
Analysis of PRDM9 substrate interactions using multi-site λ-dynamics (MSλD). *A*, Comparison of human PRDM9 and mouse Prdm9 on peptide substrates shows the highly homologous PR/SET domain have the same substrate selectivity on peptides used in this study. Graph displays mean (n=2) ±SD. *B*, Correlation of changes in relative binding free energies determined from MSλD for H3 peptides substituted at critical positions with experimental changes in methyltransferase activity for PRDM9. On-tailed Pearson (r) correlation was calculated using GraphPad Prism 9.

**Figure S3.**
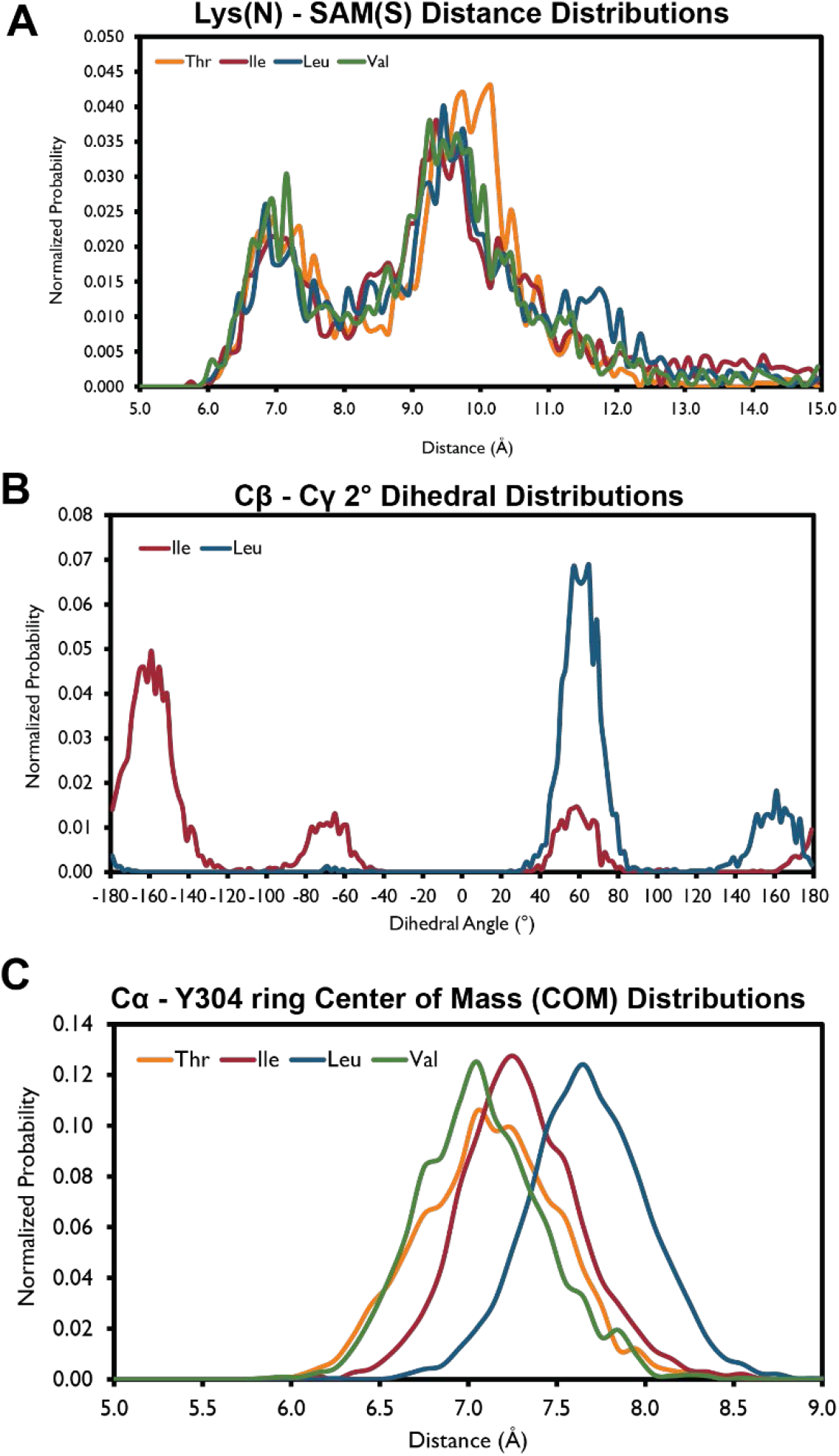
Additional MSλD trajectory analyses of PRDM9 substrate selectivity. *A*, Lys(N)-SAM(S) Distance Distributions. *B*, Cβ - Cγ 2° Dihedral Angle Distributions. *C*, Cα - Y304 ring Center of Mass (COM) Distance Distributions.

**Figure S4.**
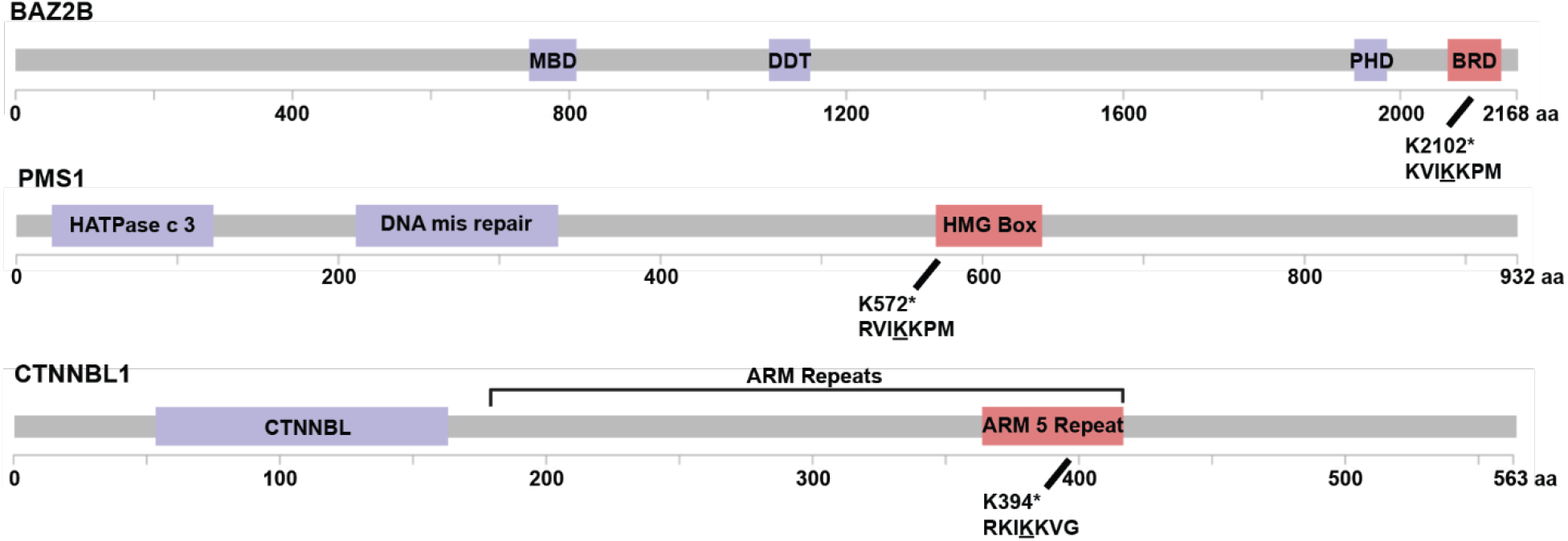
Domain maps of putative PRDM9 non-histone substrates. The predicated PRDM9 methylation site and 7-mer motif is displayed. Data for domain boundaries was derived from UniProt.

